# Modular biomaterials vaccine technology protects against multiple pathogens and septic shock

**DOI:** 10.1101/2020.02.25.964601

**Authors:** Michael Super, Edward J. Doherty, Mark J. Cartwright, Benjamin T. Seiler, Des A. White, Fernanda Langellotto, Alexander G. Stafford, Nikolaos Dimitrakakis, Mohan Karkada, Amanda R. Graveline, Caitlin L. Horgan, Kayla R. Lightbown, Frank R. Urena, Chyenne D. Yeager, Sami A. Rifai, Maxence O. Dellacherie, Aileen W. Li, Amanda R. Jiang, Vasanth Chandrasekhar, Justin M. Scott, Shanda L. Lightbown, Donald E. Ingber, David J. Mooney

## Abstract

Broad spectrum vaccines could provide a solution to the emergence of antibiotic resistant microbes, pandemics and engineered biothreat agents. Here, we describe a modular vaccine (composite infection vaccine technology (ciVAX)) which can be rapidly assembled and in which 4 of the 5 components are already approved for human use. ciVAX consists of an injectable biomaterial scaffold with factors to recruit and activate dendritic cells (DC) *in vivo* and microbeads conjugated with the broad-spectrum opsonin Fc-Mannose-binding Lectin (FcMBL) that is pre-bound to polysaccharide-rich cell wall antigens, such as the pathogen-associated molecular patterns (PAMPs) fractions, captured from whole inactivated bacteria. Vaccination of mice and rabbits with ciVAX generates potent humoral and T cell responses to PAMPs isolated from native antibiotic-resistant *E. coli* and *S. aureus*, and ciVAX protects mice and pigs against lethal *E coli* challenge in sepsis and septic shock models. In addition to the efficacy of ciVAX against homologous challenge, PAMPS isolated from an infected animal protects other animals against infection by heterologous challenge using different *E. coli* serotypes – demonstrating the potential for use of ciVAX in controlling pandemics. The advantage of the ciVAX technology is the strong immunogenicity with limited reactogenicity, the use of inactivated pathogens, and the modular manufacture using cGMP approved products which can be stockpiled ready for the next pandemic.

**One Sentence Summary:** Biomaterial vaccine induces strong immunogenicity, weak reactogenicity, and protects from *E. coli* sepsis in rodents and pigs, and MRSA skin abscess.

## Introduction

Deaths from drug-resistant microbial infections are increasing(*1, 2*), approvals of new antibiotic classes are falling(*1, 2*), and there are increasing concerns about engineered biothreat agents that may be difficult to treat with conventional therapeutics(*3*). While broad spectrum infection vaccines potentially could be used to confront challenges relating to the increase in drug-resistant bacterial infections and decrease in approvals of new antibiotics, the major antigens on bacterial cell wall surfaces are poorly immunogenic, serotype-specific polysaccharides that do not easily support vaccine production(*4, 5*). Some bacterial vaccines have been developed utilizing live attenuated bacteria (e.g. BCG for *M. tuberculosis*), but these have outgrowth risks, especially in immunosuppressed patients(*6*). Others use inactivated bacteria or inactivated toxins(*7*), but since purified carbohydrate vaccines are poorly immunogenic, and require multiple boosts(*8*), carbohydrate antigens are often chemically conjugated to proteins to increase their efficacy(*9*); however, these approaches are limited to only a subset of bacterial strains(*10*), or require modification of the antigen(*11*). Moreover, all existing vaccine modalities require that the pathogen or antigens be isolated before a vaccine can be developed, which may not be possible in a rapidly spreading pandemic or with engineered biothreat agents. Importantly, there are no approved vaccines for the most common pathogens causing sepsis, including gram negative *E. coli*, gram positive *S. aureus*, or the methicillin resistant form of *S. aureus* (MRSA).

Our goal was to develop a broad-spectrum infection vaccine that is potent, durable and safe, using complex killed bacterial components that do not require cold chain storage, and that also can be used for ring vaccination in biothreat or pandemic emergencies, without having to first identify or isolate the pathogen. To address this challenge, we combined two technologies that are currently in clinical development for other applications ― a broad spectrum engineered opsonin known as Fc-Mannose Binding Lectin (Fc-MBL)(*12-14*) that rapidly binds to carbohydrate-containing PAMP antigens (including glycoproteins and glycolipids) expressed on the surface of over 120 different pathogen species and toxins, including some that are known to trigger innate immunity (e.g., lipopolysaccharide endotoxin; LPS)(*15*), and a biomaterial scaffold-based vaccine being used for cancer immunotherapy(*16-18*) ― to create a new composite infection vaccine technology (ciVAX) (**Fig. 1**). To construct ciVAX, bacteria are inactivated using irradiation or antibiotic treatment, then the PAMPs are captured using FcMBL-coated magnetic beads (FcMBL-beads), which are mixed with mesoporous silica (MPS) microparticles with adsorbed GM-CSF and synthetic oligonucleotides rich in unmethylated cytosine and guanine nucleotides (CpG). The ciVAX is then injected subcutaneously, where the MPS particles spontaneously self-assemble into a porous biomaterial matrix, which acts as a release depot for GM-CSF that accumulates dendritic cells and CpG which activates these cells(*16*) while they process the provided antigen source. The technology is modular - to produce ciVAX to a new pathogen, all that needs to be exchanged is the PAMPS fractions from the inactivated pathogen. ciVAX can even be used for vaccination using antigens directly captured & killed from infected animals, and hence has the potential for use in autogenous biologics for veterinary applications (APHIS, USDA) as well as to rapidly address pandemics, zoonotic diseases and unknown biothreat agents.

**Fig. 1.**
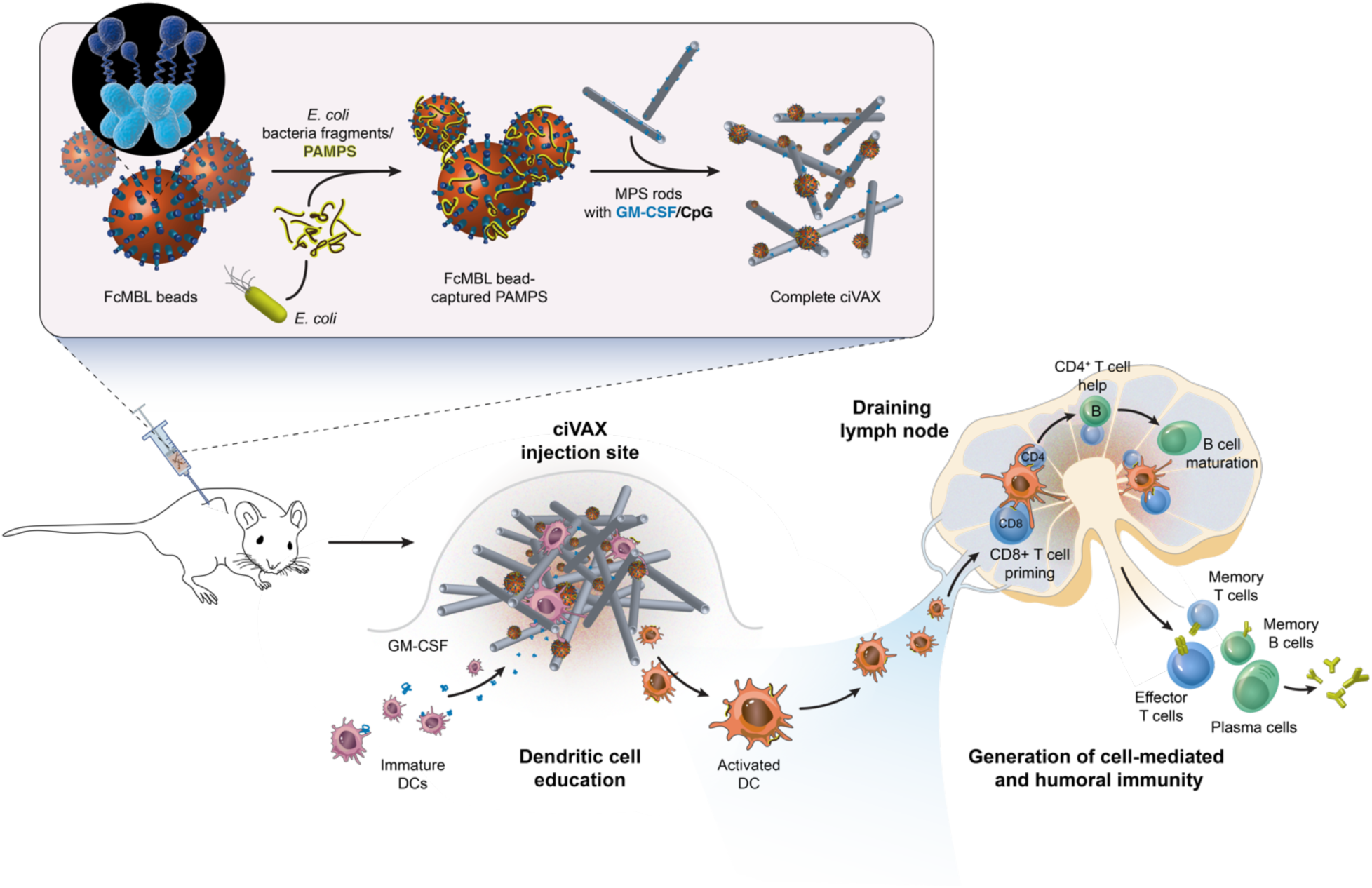
Schematic representation of production and application of the ciVAX infection vaccine for *E coli*. Fragments of killed bacteria are exposed to beads coated with the Fc-Mannose Binding Lectin (Fc-MBL), leading to capture of PAMPs on beads. The beads are then combined with mesoporous silica (MPS) rods with absorbed GM-CSF and CpG-rich oligonucleotides (CpG) to form the complete ciVAX. Needle injection of ciVAX results in accumulation of immature dendritic cells (DCs) within the scaffold formed from the MPS rods due to release of GM-CSF, and the subsequent maturation of these cells with exposure to CpG. Activated DCs traffic PAMPs from the ciVAX to the draining lymph node, interact with resident B and T cells, and generate humoral and cell-mediated immune responses to the bacterial antigens.

## Results

### ciVAX produces robust humoral and cellular immune responses with minimal reactogenicity

In our initial experiments, we did not use FcMBL to capture PAMPs and instead used the whole cell lysate made from killed RS218 *E. coli* bacteria to test immunogenicity, reactogenicity and safety in rabbits. In these pilot studies, we found that the whole cell lysate resulted in strong reactogenicity and production of necrotic lesions at the injection site (**Fig. 2A**). To avoid this toxicity, we then captured the *E. coli* PAMPs from the lysate using FcMBL-beads(*13, 14*). When FcMBL-bead-captured PAMPs containing an equivalent amount of PAMPs as the *E. coli* cell lysate (20.5 PAMP/mL vs 22.9 PAMPs/mL in lysate) was injected into rabbits (Fig. 2B), there was far less reactogenicity and little evidence of necrosis at the injection site in the FcMBL-Bead condition (**Fig. 2B**). Polyacrylamide electrophoretic analysis of FcMBL-captured PAMPs containing an equivalent amount of endotoxin LPS as the *E. coli* cell lysate (1400 and 1480 EU respectively, measured using Endosafe® LAL assay) followed by Coomassie (**Fig. 2C**) and Silver staining (**Fig. 2D**) showed that while FcMBL-bead captured material was enriched for PAMPs, it contained far less protein than the lysate. This was confirmed in MALDI-TOF Mass Spectrometry; analysis of lysate yielded multiple peptide peaks which were markedly reduced in intensity in the FcMBL-bead captured PAMPS while other peaks were enriched (Supplementary Materials Fig.1). In a head to head comparison, *E coli* RS218 lysate was added directly to MPS vaccines with GM-CSF and CpG, or PAMPs from the RS218 lysate was pre-captured with FcMBL beads first, and these were then incorporated into ciVAX. Significantly more endotoxin leached from lysate vaccines than ciVAX in a 1-hour study in vitro (**Fig. 2E**). We used the previously described sandwich ELISA-like assay(*13, 14*) to quantify and equalize the total amount of captured PAMPs added to each vaccine in all following studies to ensure equivalent PAMP dosing in different experiments, and with different bacterial species. The ciVAX forms a subcutaneous depot with little signs of necrosis, and the MPS lasts for longer than 7 days and dissolves spontaneously by 37 days (**Supplementary Materials Fig. 2**).

**Fig. 2.**
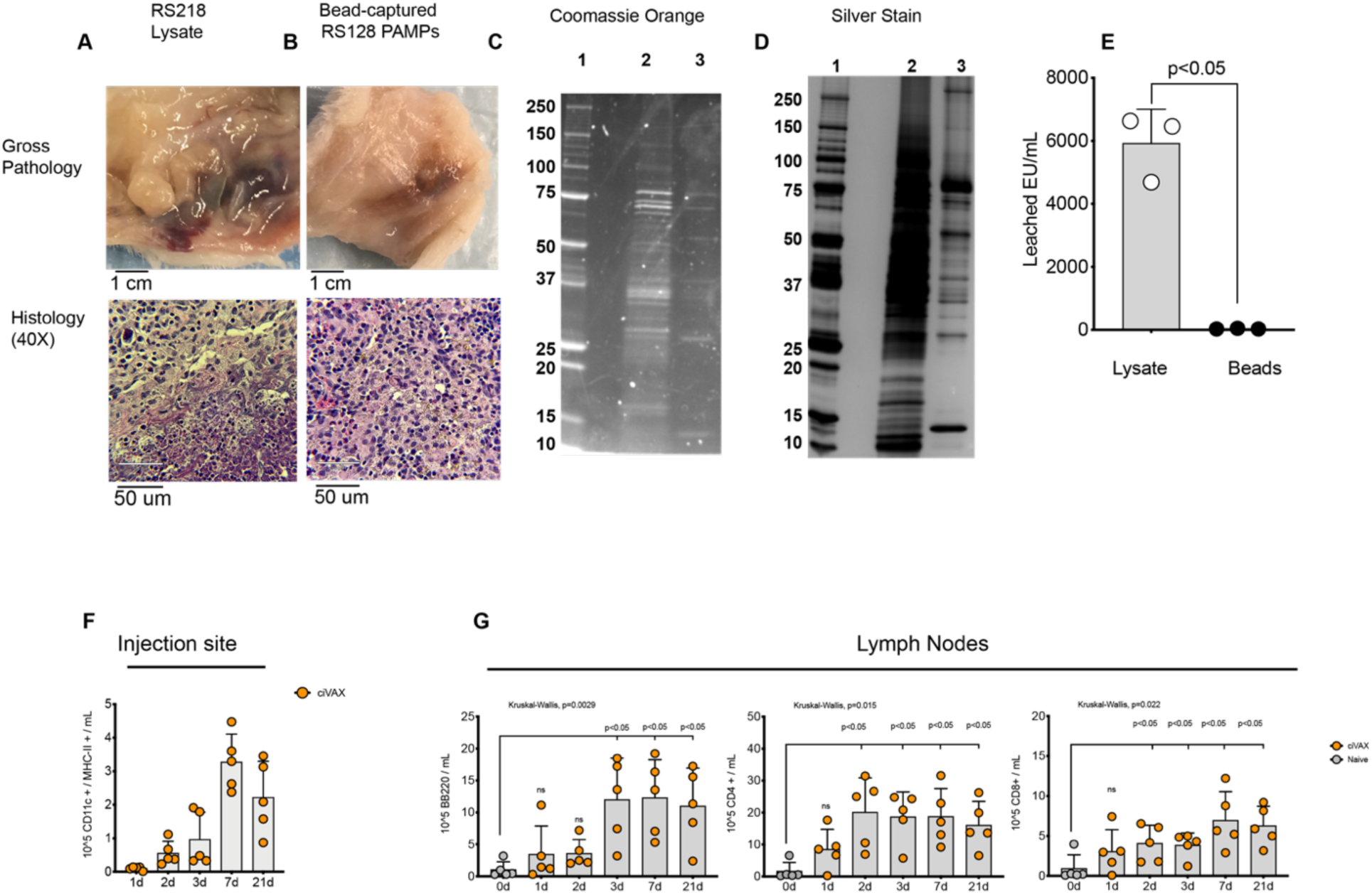
ciVAX presents a subset of bacterial cell wall PAMPs in a biomaterial scaffold that recruits DC to the injection site and produces robust humoral and cellular immune responses with minimal reactogenicity. Gross pathology and histology of injection site following whole cell lysate injection into the flank of rabbits (A) compared with sites of FcMBL captured PAMPs injection (B). Injection sites were imaged 2 weeks after immunization (n=3, one representative example shown) scale bars are 1 cm in the Gross Pathology, and 50 microns in 40 X Histology. **(C)** Coomassie Brilliant Orange stain of SDS-PAGE gel of molecular weight marker (lane 1), and equivalent amounts of endotoxin from RS218 lysate (1480 EU, lane 2) and FcMBL-bead captured PAMPs from the RS218 lysate (1400 CFU, Lane 3). **(D)** Silver stain of SDS-PAGE gel of molecular weight marker (lane 1), and equivalent amounts of endotoxin from RS218 lysate (1480 EU, lane 2) and FcMBL-bead captured PAMPs from the RS218 lysate (1400 CFU, Lane 3). **(E)** 1 hour *in vitro* study of RS218 endotoxin PAMPs leaching from MPS scaffold ciVAX without FcMBL beads (lysate) or from intact ciVAX with FcMBL beads (n=3). **(F)** Numbers of activated DCs over time, as indicated by CD11c and MHC-II staining, at the ciVAX injection site, and **(G)** numbers of B and T cells in draining lymph node, as a function of time post-vaccination (n=5 individual mice per group). Data are Mean and Standard Deviation; In **Fig. 2E and Fig. 2G** the significant differences were identified by unpaired t-test and Mann Whitney U Test respectively.

When we immunized mice with the *E. coli* PAMP ciVAX vaccine containing 7.5 ng PAMPs, we detected activated dendritic cells at the injection site within 2 days of immunization, and their numbers increased through day 7, remaining elevated through day 21 (**Fig. 2F**). This was accompanied by significantly increasing numbers of DCs, and B and T cells in the draining lymph nodes (**Fig. 2G, Supplementary Materials Fig. 3**). Past work in cancer with a similar biomaterial-based vaccine demonstrated a similar enrichment of activated dendritic cells at the injection site, and potent humoral and cellular anti-cancer immune responses.(*16, 17, 19*)

### *E coli* ciVAX propylaxis protects mice and pigs

Mice were then vaccinated and challenged with a lethal intraperitoneal injection of the homologous antibiotic-resistant RS218 *E. coli* strain (O18:H7), isolated from a human neonate with meningitis. While only 9% of the unvaccinated mice survived the challenge at 35 days, all of the vaccinated mice remained alive and healthy. Inclusion of captured PAMPs was required for protection, as survival was only 17% in sham mice receiving the same biomaterial scaffold containing GM-CSF and CpG adjuvant, but without FcMBL-bead captured *E. coli* PAMP antigens (**Fig. 3A**). Postmortem quantification of bacteria within isolated organs revealed that unvaccinated or sham-treated mice had > 1 x 10^8^ mean colony forming units (CFU) per gram in lung, liver, kidney and spleen, while vaccination reduced these numbers by 5-7 logs (**Fig. 3B**) and there was a significant increase in specific anti-RS218 antibodies in vaccinated animals (**Fig. 3C**). Moreover, the ciVAX generated a durable immune response, as a single injection protected >90 % of mice from a lethal RS218 challenge delivered 90 days after the vaccination (**Fig. 3D**) with significant decreases in bacterial counts in organs on post-mortem analysis (**Fig. 3E**) and a durable increase in anti RS218 antibody levels (**Fig. 3F**). Importantly, the infection vaccine did not significantly change the gut microbiome of rabbits over 46 days after vaccination with *E coli* or MRSA ciVAX (**Supplementary Materials Fig. 4A-C**). We next tested a homologous vaccine in a pig model of gram-negative septic shock involving intravenous injection of a human clinical isolate of *E. coli* 41949 (OM:H26), which leads to rapid renal failure and death(*20*). Animals were vaccinated and boosted (day 28) with 41949 ciVAX, and then challenged with *E. coli* 41949 on day 42. The four untreated animals developed neutropenia, thrombocytopenia, bacteremia, elevated lactate, acute kidney injury with extremely high creatinine levels, and porcine systemic organ failure (SOFA)(*20*), indicating fulminant sepsis within 12 hours, which required that they be euthanized (**Fig. 3G**). In contrast, the four vaccinated pigs had increased neutrophil counts, low lactate levels, normal SOFA scores, creatinine levels below the euthanasia threshold (**Fig. 3H-I** and **Supplementary Materials Fig. 5**), and they all survived to the end of the experiments at 28 (n=2) and 72 hours (n=2) respectively (**Fig. 3G**). Importantly, the vaccinated pigs were clinically healthy, despite live pathogens and dead pathogen-derived PAMPs remaining in their bloodstream (**Supplementary Materials Fig. 5**), suggesting that the ciVAX vaccine protected the pigs from endotoxic shock through a strengthened humoral and cellular immune response, rather than a bactericidal mechanism.

**Fig. 3.**
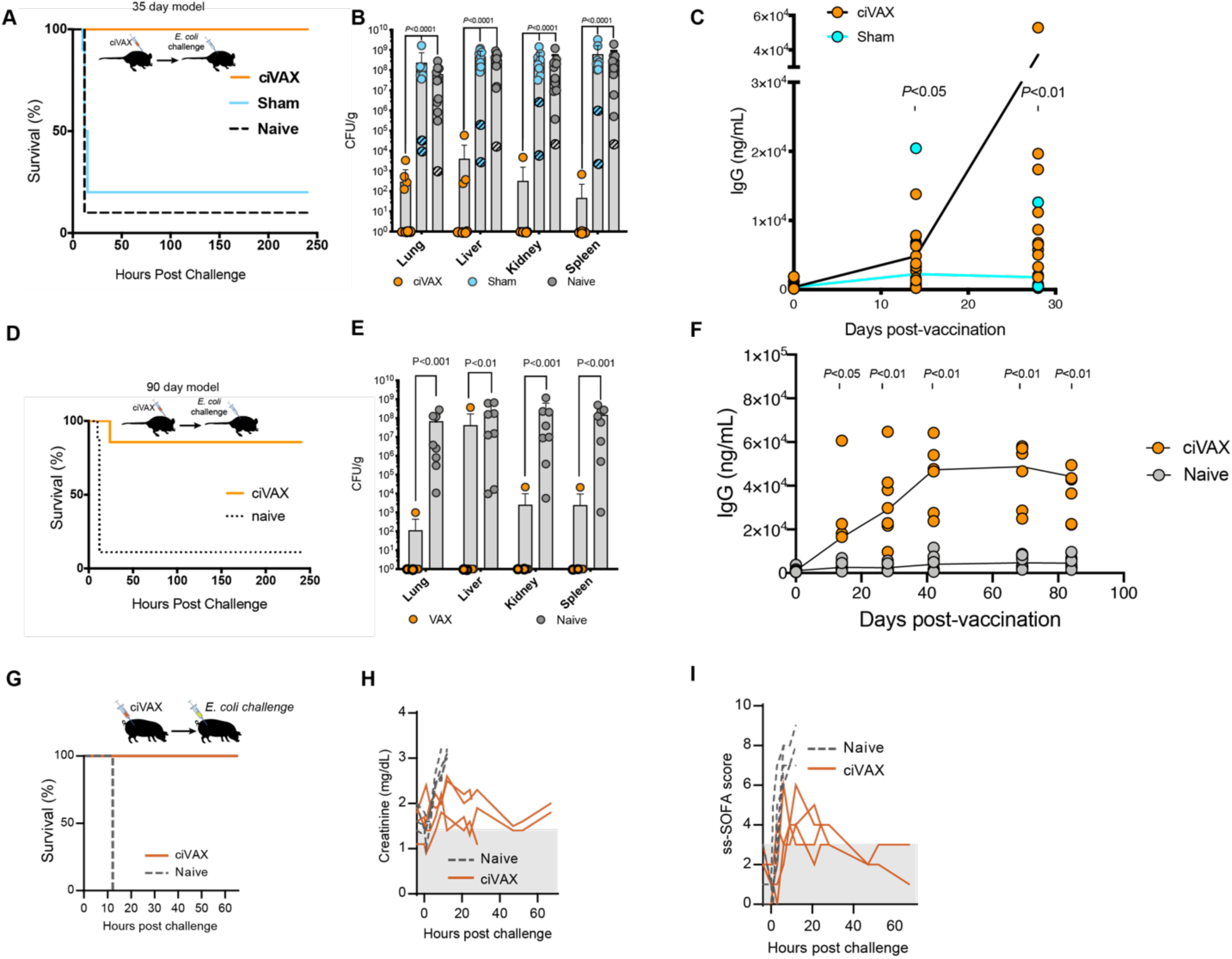
ciVAX protects mice from *E. coli* sepsis in an RS218 challenge model, and pigs from *E. coli* septic shock. (**A-F)** ciVAX was produced by mixing lyophilized MPS biomaterial scaffold containing murine GM-CSF and CpG with cefepime-killed RS218 captured on FcMBL paramagnetic MyOne Beads. 8-10 week old Male and female BALB/c mice were vaccinated with a single subcutaneous injection with ciVAX (orange line), or sham scaffold with GM-CSG and CpG but without the FcMBL/RS218 (blue), or unvaccinated control (gray dashed). (**A-C)** Mice were challenged at 35 days (10-12 per group) or (**D-F)** at 90 days (8-9 per group) post-vaccination with an intra-peritoneal injection of 5e6 CFU *E*.*coli* (RS218). Mice were monitored for survival for a further 240 hours (10 days). At humane criteria endpoints or at the conclusion of the experiment, mice were sacrificed and organs were disaggregated using bead-milling followed by quantitative culture. (**B**,**E**) Quantification of bacteria from isolated organs in the various experimental groups. Hatched blue and gray symbols depict the sham and naïve mice that survived the challenge. (**C**,**F**) RS218-specific antibodies in mouse serum were measured after vaccination and before challenge, using F(ab’)2 Fragment Goat anti mouse IgG (Jackson Labs). **(G-I)** ciVAX for swine was prepared as above, using multidrug-resistant clinical isolate *E. coli* 41949 (BWH Biorepository) instead of RS218. Female swine (4 per group) were vaccinated, boosted at d28 and challenged 42 days after vaccination with a lethal dose of 41949 delivered by infusion over 4-8 hours using the conscious model of sepsis we developed (*20*). Yorkshire pigs were analyzed over time for survival (**G**), creatinine levels (**H**), and Swine-specific Sequential (Sepsis-related) Organ Failure Assessment (ssSOFA) (**I**) in both vaccinated (orange) and naïve (dashed) swine. **Figs. (3B), (3E):** Data are mean and standard deviation; **Figs. (3B-F):** statistically significant differences between the independent groups were defined by non-parametric Mann-Whitney *U* test.

### ciVAX can be based on other biomaterial scaffolds and requires all agents

While the studies described to this point were performed with one biomaterial vaccine system (MPS rods), this general approach is likely to be useful with a variety of material-based vaccine strategies. To test this possibility, PAMPs bound to FcMBL-beads were combined with another biomaterial-based vaccine currently in a clinical trial for the treatment of Stage IV melanoma (WDVAX; NCT01753089). Vaccination with PLG ciVAX containing cefepime-killed RS218 captured on FcMBL beads promoted survival in a high fraction of mice subsequently challenged 35 days post-vaccination (**Fig. 4A**), and dramatically reduced bacterial titers in all organs analyzed **(Fig. 4B**). Vaccines fabricated without GM-CSF/CpG or PAMP/FcMBL, or missing all actives provided no survival benefit and led to high bacterial load in organs of challenged mice (**Fig. 4A-B**). These findings indicate that both the antigen accumulation/activation function of the scaffold component of ciVAX, and the antigen presentation via FcMBL is required for the efficacy of these vaccines. Mice challenged 90 days post-vaccination with the PLG ciVAX also demonstrated high survival (**Fig. 4C**), supporting the durability of the induced immune responses. As the PLG-based ciVAX, unlike the MPS rods, requires surgical implantation, all subsequent studies were performed with the MPS-based ciVAX.

**Fig. 4.**
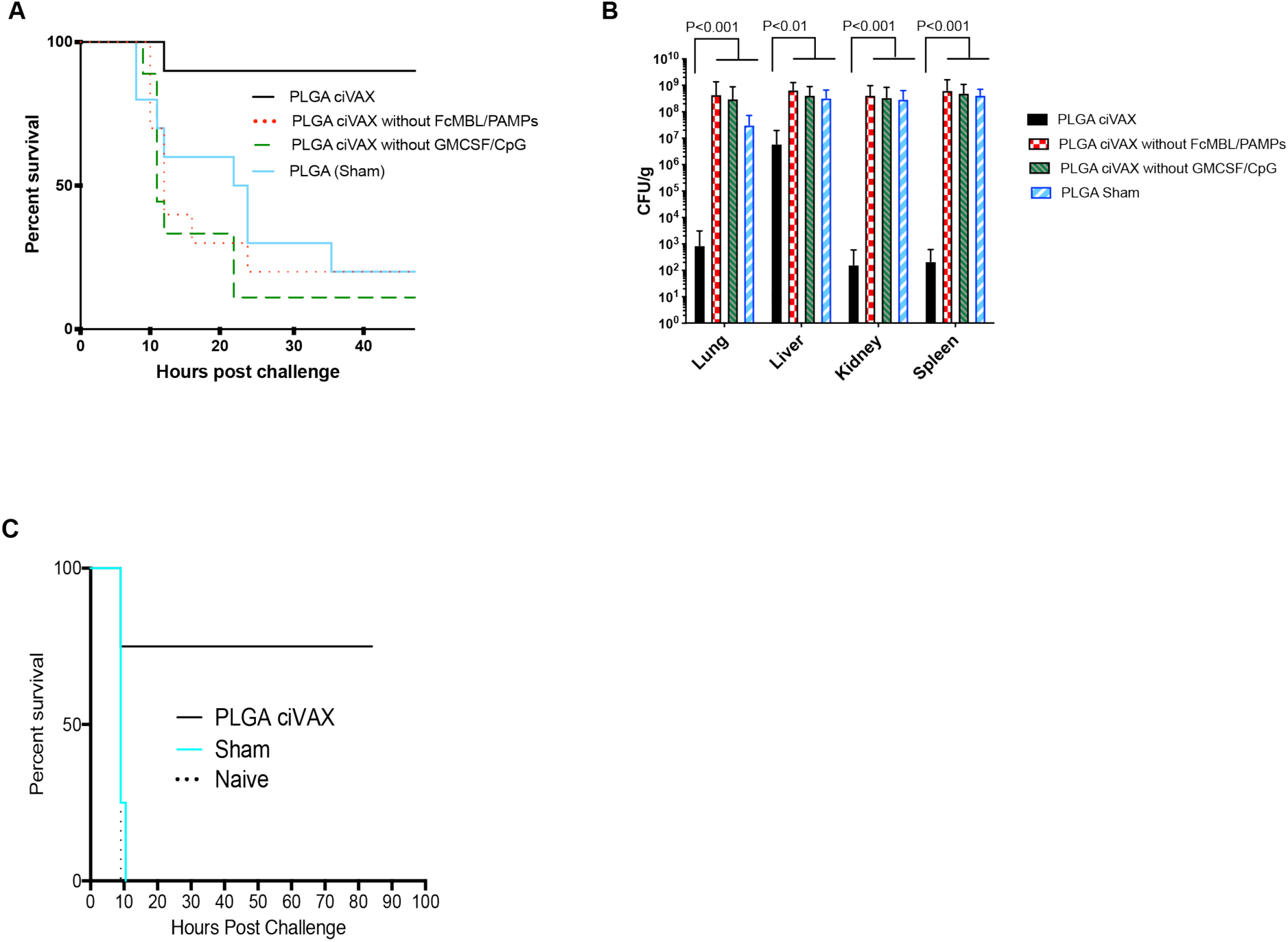
PLGA ciVAX vaccine is effective compared with individual components and yields a durable, protective, immune response. **(A-C)**, PLGA ciVAX was produced by mixing PLGA scaffold containing murine GM-CSF and CpG with cefepime-killed RS218 captured on FcMBL myOne beads followed by lyophilization.**(A)** 8-10 week old male and female mice were vaccinated with a single subcutaneous implantation of the ciVAX (black line), or ciVAX without the FcMBL/PAMPs (red dotted line), or ciVAX without the GMCSF/CPG (green dashed line) or just PLGA sham (blue line) (n=5 per group). Mice were challenged at 35 days post-vaccination with an intra-peritoneal injection of 5e6 CFU *E*.*coli* (RS218) and monitored for mortality and humane criteria for a further 48 hours. **(B)** Pathogen load in organs at necropsy for the ciVAX group and the groups without various components of the vaccine (n=9 per group). **(C)** 8-10 week old male and female mice were vaccinated with a single subcutaneous implantation of the ciVAX (black line), or ciVAX sham without the FcMBL/PAMPs (blue line), or unvaccinated (naïve) (dotted red line) (n=5 per group) followed by challenge with 5e6 CFU *E*.*coli* (RS218) at 90 days after vaccination. Data are mean and standard deviation; Statistically significant differences between multiple independent groups were identified by implementing non-parametric Kruskal-Wallis test.

#### *S aureus* ciVAX propylaxis reduces skin abscesses in mouse model

*S. aureus* is the most common gram-positive organism causing sepsis and many *S. aureus* strains are antibiotic resistant, yet there is no approved vaccine for this pathogen(*21*). Mice immunized with ciVAX containing PAMPs isolated from MRSA mounted potent, durable, and specific IgG responses that were multiple fold higher than in sham animals (**Fig. 5A**). Vaccinated mice that were subsequently challenged with MRSA infection demonstrated significantly fewer skin abscesses compared to controls (**Fig. 5B**).

**Fig. 5.**
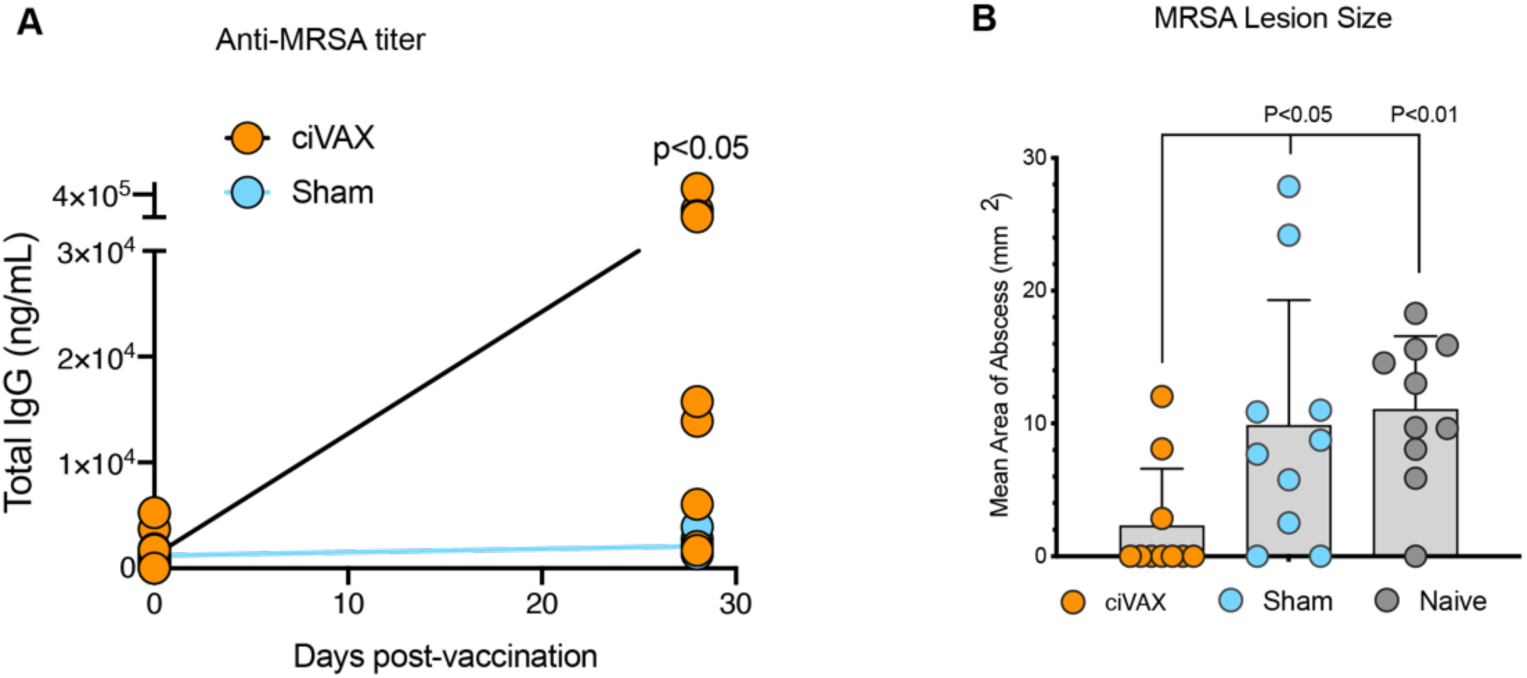
ciVAX provides functional immune protection to gram positive *S. aureus*. ciVAX was produced by mixing lyophilized MPS biomaterial scaffold containing murine GMCSF and CpG with vancomycin & daptomycin-killed MRSA (ATCC 12598) captured on FcMBL paramagnetic MyOne Beads. **(A)** 8-10 week old female BALB/c mice were vaccinated & boosted (day 14) by subcutaneous injection with ciVAX (orange), or sham scaffold without the FcMBL/RS218 (blue), or unvaccinated control (gray) (n=10 per group). MRSA-specific antibodies in mouse serum were measured using vancomycin & daptomycin-killed MRSA adsorbed directly on well-plates, blocked to reduce interaction between Staphylococcal protein A and IgG. **(B)** Mice were challenged 35 days after vaccination with 5e7 CFU MRSA on the left flank and monitored for abscess formation for a further 72 hours. Data are mean and standard deviation; statistically significant differences between the independent groups were identified by non-parametric Mann-Whitney *U* test.

### Potential for ciVAX use in pandemic applications

The broad-spectrum binding capabilities of FcMBL could enable development of vaccines from PAMPs extracted from blood or tissues of infected individuals. In veterinary medicine these could qualify as Autogenous Biologics for approval by USDA-APHIS (9 CFR 113.113). In one world health and pandemic applications, the ciVAX technology could be used in ring vaccination to protect others in the community from infection by the original pathogen or even by closely related species and variants, potentially providing benefit even where there is insufficient time for full identification of the organism. As it was impractical to prepare ciVAX from mice due to their limited blood volumes, to model ring vaccination, PAMPs were magnetically captured in less than one hour from the blood of a pig previously infected with *E. coli* 41949 (OM:H26), using FcMBL-beads, which were then combined with the lyophilized biomaterial components. After this single heterologous vaccination with captured *E coli* OM:H26 PAMPs, we challenged the mice with a lethal dose of a different *E coli* serotype (O18:H7). All unvaccinated mice died following *E. coli* RS218 (O18:H7) challenge, and organ bacterial counts were > 1 x 10^8^ CFU/g (**Fig. 6A-B**). In contrast, 80% of the vaccinated mice survived the lethal *E. coli* RS218 challenge, and there was a significant decrease in pathogen load in lung and kidney at the end of the 28 day study, and trends for decreased pathogen in the spleen and lungs (**Fig. 6A-B**).

**Fig. 6.**
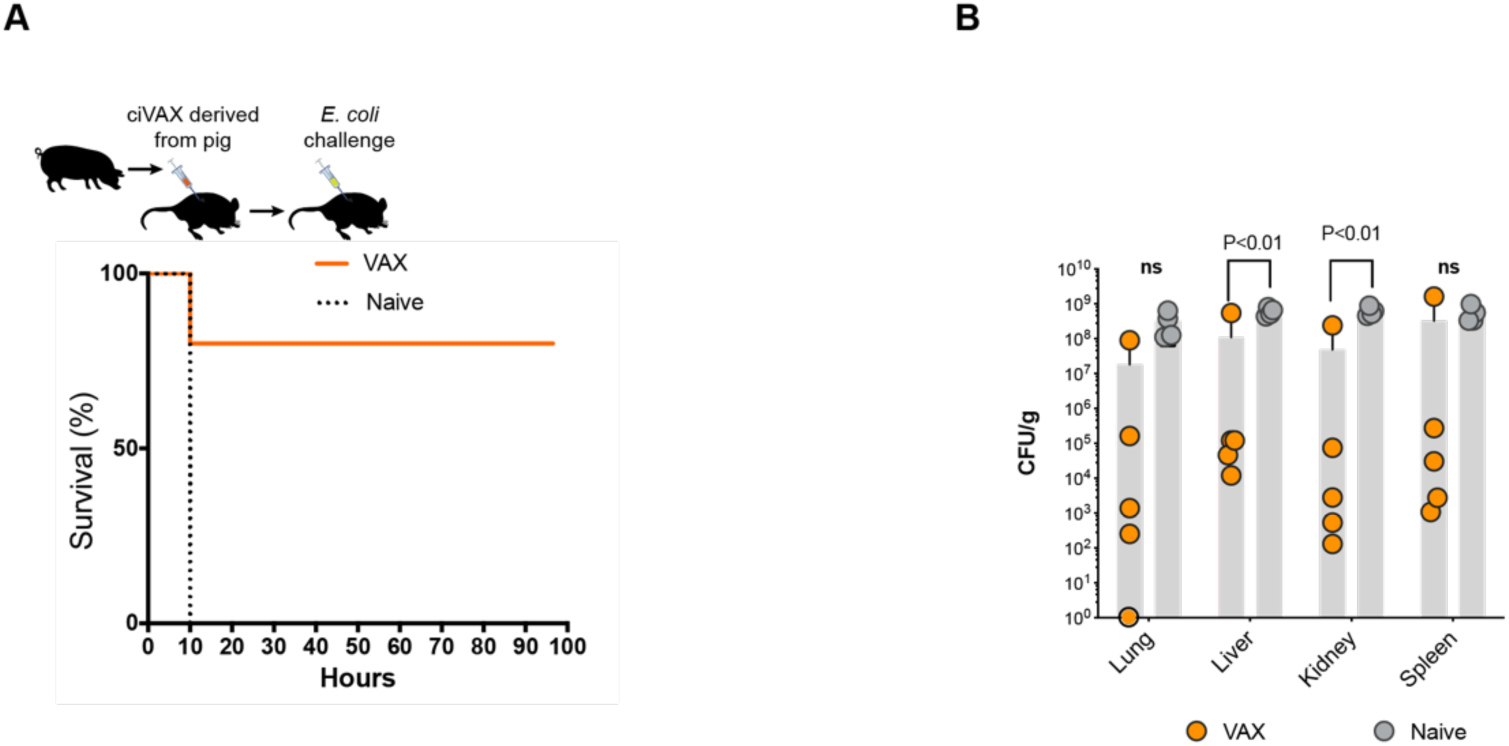
ciVAX produced from an *E. coli* infected animal could be used to protect against another *E. coli* serotype. ciVAX was produced by capturing *E. coli* 41949 (OM:H26) PAMPs directly from the blood of an infected pig treated with Cefepime, using FcMBL-coated superparamagnetic beads, and then mixed with lyophilized MPS biomaterial scaffold containing murine GMCSF and CpG. (**A**)Five mice were vaccinated (orange line) and at d35 these mice (and 5 naïve mice controls – dotted line) were challenged with lethal doses of *E. coli* RS218 (O18:H7) and followed for 96 hours. (**B**) At humane endpoints or the end of the experiment, mice were sacrificed and the pathogens in lung, liver, spleen and kidney were measured by quantitative culture on sheep blood agar. Data are mean and standard deviation; significant differences were identified by unpaired t-test.

## Discussion

The ciVAX technology represents a potentially significant advance in the field of infection vaccines, as there are no approved vaccines for the most common pathogens causing sepsis. A four-valent vaccine comprising *E. coli* O1A, O2, O6 and O25B antigens conjugated to *Pseudomonas aeruginosa* exoprotein A is in Phase 2 trials for prevention of extra-intestinal *E. coli* (ExPEC) infections(*22*). While this vaccine led to an increase in functional antibodies to at least 2 of the 4 antigens, it is estimated that more than 10 different antigens will be needed to cover more than 60% of ExPEC and 90% of meningitis isolates. There are no approved prophylactic or therapeutic vaccines for *S. aureus* or MRSA infection. There have been attempts to develop vaccines to *S. aureus* polysaccharide type 5 or 8 conjugates, but these have not been successful in clinical trials and newer trials are again focused on increasing the number of antigens(*21*). We anticipate ciVAX could be more effective than these approaches because multiple antigens (including glycoproteins and glycolipids) are included in their native forms, antigens are presented on particulate substrates and slowly released from biomaterial depots, yielding strong immunogenicity with little toxicity and reactogenicity, and ciVAX includes bioactive molecules that recruit large numbers of dendritic cells for *in vivo* priming and maturation, which promote development of potent affinity-matured immune responses. The benefit of ciVAX containing multiple (polyclonal) antigens, rather than a sub-set of immunodominant antigens expressed using recombinant DNA technology, may also be a limitation as the PAMPs material is not defined and all the antigens have not been identified. It is possible that ciVAX could contain antigens causing immunosuppressive reactions rather than protective immune responses. Further, the impact of cellular immunity versus humoral immunity needs to be determined, and further studies are needed to determine how ciVAX will work in those with immature or weak immune systems. It will also be important to explore therapeutic vaccinations of this technology to expand its potential clinical utility.

In summary, this modular ciVAX PAMP vaccination technology can be used to mount strong immune responses to Gram negative and Gram positive bacteria following subcutaneous administration. The system produced life-saving effects in small and large animal sepsis models resulting from infection with *E. coli* from different serotypes. The broad-spectrum efficacy against MRSA and multidrug-resistant bacteria suggest that this technology could represent a new alterative to antibiotics in the post-antibiotic era. Importantly, given the broad spectrum PAMPs binding capabilities of FcMBL(*14*), this vaccine approach should also be effective against a wide range of fungal, parasitic and viral targets. Moreover, the modular nature of the technology, and the use of lyophilized components should make this vaccination particularly useful for global and one world health applications. The ability to target PAMPs from a broad spectrum of pathogens without time-consuming culture steps also raises the possibility that it could provide a unique clinical countermeasure to quickly control spread of infection in pandemics and biothreat scenarios. In this situation the entire ciVAX except the antigen could be prefabricated from GMP sourced components and stored in lyophilized form at strategically located depots. On obtaining the pathogen, it could be inactivated, captured on lyophilized FcMBL microbeads and rapidly added to the other lyophilized components. We have documented this assembly is possible in a 1-hour timescale.

## Materials and Methods

Preparation of bacterial antigens. *S. aureus* 12598 (ATCC 12598), *E. coli* RS218 (kindly provided by James R. Johnson from the University of Minnesota), *E. coli* 41949 (Brigham and Women’s Crimson Biorepository), were grown individually in RPMI 10mM glucose at 37°C 225rpm to a McFarland of 1.0 (∼2 × 10^8^ CFU/mL). The cultures were placed on ice for 10 minutes to slow growth then centrifuged at 3,000 × g for 10 minutes at 4°C (Eppendorf 5810R). The supernatants were discarded and the pellets were resuspended in fresh RPMI 10mM glucose until a McFarland of 5.0 (∼10^9^ CFU/mL) was achieved. Next, 1mg/mL of cefepime (NDC 25021-121-20) was added to *E. coli*, 1 mg/mL vancomycin (NDC 0409-4332-49) and 0.5 mg/mL daptomycin (Toronto Research Chemicals, Canada) was added to *S. aureus*. All cultures were then placed back in the incubator overnight (∼17 hours) at 37°C 225rpm. The following morning, each culture was passed through a 70 µm cell strainer (Sigma-Aldrich, USA) to remove large cellular debris. A second dose of the respective antibiotic was added to each culture and incubated overnight (∼17 hours) at 37°C 225rpm. Each culture was then plated in triplicate on TSA II Soy Agar with 5% Sheep Blood (Becton Dickinson, USA) and placed in 37°C 5% CO_2_ incubator for 24 hr to confirm complete bacterial death by absence of CFU. Afterwards, the cultures were stored a −80°C. When ready, the cultures were thawed and diluted 1:2 in 50mM Tris-HCl, 150mM NaCl, 0.05% Tween-20, 5mM CaCl_2_, pH 7.4 (TBST 5mM CaCl_2_) (Boston BioProducts, USA). FcMBL beads^13^ (Biotinylated FcMBL on 1µM Dyna MyOne Dynabead [Thermo Fisher Scientific, USA]) were added at 25µl/mL of a 5 mg/mL stock, and the solution was mixed at room temperature for 30 minutes using a hula mixer (Thermo Fisher Scientific, USA) set to 20 rpm. After FcMBL capture/enrichment of bacterial PAMPs, the solution was magnetized (Thermo Fisher Scientific, USA) for 10 minutes to allow removal of supernatant without bead disruption. The tubes were removed from the magnet and fresh TBST 5mM CaCl_2_ was added back at equal volume to wash the beads. Finally, the tubes are inverted until the beads were completely homogenized. This solution was stored at −80°C until needed.

### Quantification of bacterial antigens captured on FcMBL beads

The captured PAMPs bacterial antigens captured on FcMBL beads were quantified using an Enzyme-Linked Lectin Sorbent Assay (ELLecSA) as previously published (*13*). Briefly, FcMBL was biotinylated at the N terminus of the Fc protein using an N-terminal amino-oxy reaction and then coupled to streptavidin superparamagnetic beads (1µM MyOne Dynabead (Thermo Fisher Scientific, USA). Samples were screened using 5 µg of the FcMBL beads, 200 µL test sample, and 800µL TBST 5mM CaCl_2_ supplemented with 10mM Glucose, incubated for 20 minutes at 22°C 950 rpm in a plate shaker (Eppendorf, USA). The automated magnetic-handling system (KingFisher™ Flex (Thermo Fisher Scientific, USA)), was used for washing steps followed by colorimetric development with horseradish peroxidase (HRP)-labeled MBL. The ELISA was quenched using sulfuric acid and read at OD 450 nm and compared with a standard curve of the well-known yeast PAMPs - Mannan.

### Preparation of vaccine scaffolds

#### Mesoporous silica rods (MPS)

To synthesize high-aspect-ratio MPS (88 μm × 4.5 μm), 4 g of P123 (Sigma-Aldrich, USA) surfactant (average M_n_ ∼5,800) was dissolved in 150 ml of 1.6 M HCl solution then stirred with 8.6 g of tetraethyl orthosilicate (TEOS, 98%, Sigma-Aldrich, USA) at 40 °C for 20 h, followed by incubation at 100°C for 24 h. To extract the surfactant, the as-synthesized particles were refluxed for 10 h in 1% HCl in ethanol. The resulting MPS particles were then filtered, washed with ethanol and dried. MPS morphology was measured using optical microscopy. To make the completed MPS vaccine, 3 µl GM-CSF (Murine Gm-CSF, Peprotech, USA; Human - Leukine® for pig and rabbit, Sanofi, USA) and 100 µl CpG (Murine CpG, ODN 1826, Human CpG ODN 2006 for pig and rabbit, InvivoGen, USA) were both added at 1mg/ml in WFI. Captured antigen, defined antigen, or lysate (e.g. 25 µl of FcMBL beads with captured 15 PAMPs units of enriched *E coli* lysate) were then added to the 10 mg MPS solution. The resulting slurry is sonicated for 5-10 seconds in a water bath and mixed overnight at room temperature using a HulaMixer (Thermo Fischer Scientific, USA) at 45 rpm. The mixture was then frozen at -80°C for a minimum of 12 h prior to lyophilization. Lyophilization is for a minimum of 48 hours. The material is reconstituted in WFI prior to injection.

#### Poly(lactic-co-glycolic acid) (PLGA)

18 mg PLGA microspheres (Phosphorex, USA) containing 9 µg of encapsulated GM-CSF (Murine GM-CSF, Sanofi, USA; Human - Leukine® for pig and rabbit, Peprotech, USA) were combined with a 1 mL solution of captured antigen, defined antigen, or lysate (e.g. 25 µl of FcMBL beads with captured 15 PAMPs units of enriched *E coli* lysate) vortexed for 5 minutes before being flash frozen with LN_2_ and lyophilized for 72 – 96 hours. The lyophilized powder was combined with 300 µg of CpG (Murine CpG, ODN 1826, Porcine, Rabbit, Human CpG ODN 2006 for pig and rabbit, InvivoGen, USA) which has been condensed with Polyethyleneimine (PEI) and 130 mg of sieved sucrose (250 - 450 µm) then manually mixed until homogenous. This homogenous powder was poured into a die set and formed using a Carver press under 1500 psi of pressure for 45 seconds, resulting in a solid disc of approximately 2 mm high by 9 mm diameter. The disc was placed in a pressure chamber (Parr, USA) overnight and exposed to 800 psi CO_2_ (∼18 hr). Prior to implantation, the disk is placed in 10 ml water for injection (WFI) (Thermo Fisher Scientific, USA) which dissolves the sieved sucrose, creating the final porous structure for implantation. This step reduces the vaccine dose by approximately 50-60%.

## Animal Models

### Mice

All studies were performed in accordance with institutional guidelines and approved by the Institutional Care and Use Committee of Harvard Medical School. Male and female BALB/c mice (8-10 weeks old, weighing 18-25g, Charles River Labs) were used in the experiments and housed in a BL2 room under standard conditions. Animals were provided autoclaved bedding, enviro-dri nesting material, had free access to food and water, and were kept on a 12 h dark: 12 h light cycle.

### Vaccination

Mice were briefly anesthetized, fur clipped, and 250uL of vaccine (vortexed before injection) placed subcutaneously on the dorsum or flank. Vaccine sites were monitored over time for irregularities. Serum was collected from mice at 14 day intervals. Mice vaccinated with ciVAX MRSA were boosted at d14 with ciVAX MRSA.

### RS218 and MRSA Challenges

Male and female BALB/c mice were challenged with 5e6 CFU *E. coli* (RS218) IP in 100uL. Animals were monitored several times per day for mortality and humane criteria. Animals presenting as moribund were euthanized and were considered deceased. Female BALB/c mice were challenged with 5e7 CFU MRSA on the left flank at day 35. Animals were briefly anesthetized, fur clipped, depilatory cream applied, and the pathogen placed subcutaneously in 50uL. Animals were monitored daily for lesion size and progression. Sustained release buprenorphine and Extended Data fluids were provided for all challenge models. At termination organs were extracted for immunological and CFU quantification.

### Rabbits

Approval for the study was obtained from the Institution Animal Care and Use Committee (PARF). *E. coli* RS218 was cultured in RPMI 10 mM glucose media at 37°C 220 rpm until a McFarland of 0.5 (1 × 10^8^ CFU/ml) was achieved. The culture was then spun at 3000*g* for 15 min at 4°C to pellet the bacteria. The pellet was resuspended using RPMI 10 mM glucose until a McFarland of 5.0 (1 × 10^9^ CFU/ml) was achieved. The antibiotic, Ceftriaxone (NDC 60505-6104-4), was added at 1 mg/ml and the culture was incubated overnight at 37°C 220 rpm. The following day, cellular debris was removed by filtering through a 70 µm membrane, and the culture was treated again with 1 mg/ml Ceftriaxone overnight. Afterwards, the culture was plated in duplicate on Trypticase Soy Agar II 5% Sheep Blood to ensure complete bacterial death confirmed by absence of CFU after incubation at 37°C overnight. To make the 50% lysate solution, the culture was diluted 1:2 in TBST 5mM CaCl_2_. To make the 1:2 bead captured solution, the culture was diluted 1:2 in TBST 5mM CaCl_2_ and 1 µm FcMBL beads were added at a concentration of 25 µL/ml. This solution was then incubated at room temperature for 30 min using end-over-end mixing. Beads with captured PAMPs are magnetized for 5 min and the supernatant was removed. An equal volume of TBST 5mM CaCl_2_ was added back, and the beads were inverted to homogenize in solution. 3 rabbits were injected by subcutaneous route using either the 50% whole cell lysate or the 1:2 bead captured solution containing equivalent PAMPs levels (22.9 PAMPs/ml ±1.9 and 20.5 PAMPs/ml ±4.2 respectively). Rabbits were boosted at day 28 with the same preparations used for priming. At the conclusion of the study at day 42, the vaccination sites were compared by gross pathology and histology.

### Pigs

Approval for the study was obtained from the Institution Animal Care and Use Committee (protocol number 17-03-3400). The work was conducted at an AAALAC accredited and USDA registered facility and in accordance with NIH guidelines. Outbred juvenile Yorkshire swine (40-50kg) were vaccinated against *E. coli* 41949 via two subcutaneous injections of four milliliters in the left or right flank. Initial vaccination occurred 43 days prior to challenge and was 16 times the mouse dose. The animals received a booster vaccine on day 27/28. The booster was given as a single injection of four milliliters subcutaneously on the left flank and was four times the mouse dose.

On Day 43, animals were anesthetized with intramuscular injections of atropine (0.04 mg/kg), telazol (4.4 mg/kg) and xyalzine (2.2 mg/kg) and maintained on isoflurane (1.5–2.0%) and oxygen (0.5-2.0 liter/min) delivered through an 8.5-9-mm endotracheal tube using a positive pressure ventilator. Animals were placed in the supine position and a 20-g intravenous cannula was placed in the left or right marginal ear vein for administration of drugs and fluids, and sterile saline (0.9% NaCl) was administered continuously at a rate of 200-300 ml/hr. A 10-12-Fr Foley catheter (SurgiVet, Smiths Medical) was placed for urinary drainage, and a 5 to 6 Fr percutaneous sheath catheter (Arrow, Teleflex, NC) was placed in the left or right femoral vein for obtaining a blood sample before surgery to serve as a baseline and for pathogen administration after surgery. Ventilator settings were determined on an individual basis depending on the pig size.

Bilateral neck cutdowns were performed to expose and isolate the right carotid artery and left and right jugular veins. The right carotid and right jugular were cannulated with 8 Fr sheath catheter (Arrow). In two animals, the left jugular was cannulated with a 7 Fr triple lumen catheter (Arrow) to allow for interventions if needed. After cannulations were complete, the incisions were closed. The animals remained under anesthesia and a bolus of 1.2-1.5 × 10^5^ CFU/kg E. coli 41949 (clinical isolate 41949 obtained from Brigham and Women’s Hospital, Crimson Biorepository, USA) was administered via the femoral vein at a rate of 1ml/min over 10 minutes using a syringe pump. Following the bolus, the femoral line was pulled, the animals were recovered from anesthesia and returned to housing. Once recovered, the animals received a second infusion of 3.51 × 10^7^ - 1.36 × 10^8^ CFU/kg *E. coli* 41949 via the right jugular vein over 4-7 hours using a syringe pump.

Animal vitals were monitored, and blood samples were taken for the following analysis: blood culture, complete blood count, blood gas, blood chemistry, clotting profile, and enzyme linked lectin-sorbent assay. Organ samples were obtained at the conclusion of the study.

### Bacterial organ culture

#### Mous

Lung, liver, kidney, and spleen tissue were collected and placed in 2 mL sterile tubes containing 2.38 mm metal beads (MO BIO/QIAGEN, USA). 1 mL of sterile saline is added to each tube, weighted, then fragmented by bead mill treatment at 30 Hz for 3 min using a Mixer Mill MM 400 machine (Verder Scientific, Inc., USA). Bacterial CFUs were quantified as CFU/g from the resulting slurry by spiral plating (Eddy Jet 2, IUL, Spain), automated counting (Flash & Go, IUL, Spain) and adjusting for the volume of water added to organs.

#### Pig

To determine the CFU/g of organs, two 0.5-1 cm3 samples from 4 distinct locations per organ (lung, liver, kidney, spleen) were placed into separate pre-sterilized Nalgene bottles (Thermo Fisher Scientific, USA) containing stainless steel balls (McMaster-Carr, USA) and weighed. 17 ml of sterile water was added to each. Organ tissue was fragmented by bead mill treatment at 30 Hz for 3 min using a Mixer Mill MM 400 machine (Verder Scientific, Inc., USA). CFUs were quantified as *E. coli* CFU/g from the resulting slurry by spiral plating (Eddy Jet 2, IUL, Spain), automated counting (Flash & Go, IUL, Spain) and adjusting for the volume of water added.

### ELISA for IgG Antibody titers to RS218 and MRSA

For RS218, Polystyrene microwell plates were functionalized (High-Bind ELISA plate, 96 well, Santa Cruz Biotechnology Cat# sc-204463) with Mannose Binding Lectin, by adding 100 μL of a 0.5 μg/mL MBL (Sino Biological, Inc., China) in TBST 5mM CaCl_2_ (Boston BioProducts, USA). Plates were blocked overnight with 1% BSA. For MRSA the Polystyrene microwell plates were directly coated with 100μL of MRSA antigens. MRSA plates were additionally incubated in 5% pig serum in 1% BSA in TBST and blocked for 1 hour and 30 min at room temperature. For both ELISAs, serum samples were diluted at 1/100, 1/1000 and 1/10000 in blocking buffer (1% BSA in TBST) and stored at 4 °C until used. Subsequently, the wells were washed six times and samples were added to the wells and incubated for 1 h and 30 minutes at room temperature. After six washes, Peroxidase-AffiniPure F(ab’)2 Fragment Goat anti mouse IgG (Jackson ImmunoResearch #111-036-047) diluted to 1/20,000 was added to the wells and incubated for 1 hour at room temperature. The wells were washed six times and 100μL of TMB (3,3’,5,5’-tetramethylbenzidine) detecting HRP buffer was added. After 15 min the color development was stopped with 1 M H2SO4. Optical density (OD) was measured at 370 - 652 nm using Microplate Reader and the data were processed using GraphPad Prism software. The samples were run in duplicates unless otherwise stated and standards were run in triplicates

### Flow Cytometry

To evaluate the phenotype of cells isolated from the injection site and lymph nodes, we used panels of fluorescent conjugated monoclonal antibodies as well as additional isotype controls. Cells were stained with APC-conjugated anti-mouse BB220 in combination with FITC-conjugated anti-mouse GL7, APC/Cy7-conjugated anti-mouse CD3, BV421-conjugated anti-mouse CD4, BV711-labelled anti-mouse CD8 for detecting lymphocytes and PE-labelled anti-mouse CD86, PE/Cy7-labelled anti-mouse CD11b, FITC-labelled anti-mouse CD11c, APC/Cy7-conjugated anti-mouse I-A/I-E and BV421-conjugated anti-mouse CD80 monoclonal antibodies. Nonspecific staining was avoided by adding an FcR Blocking Reagent and LIVE/DEAD Fixable Dead Cell stain. Cells were incubated for 30 minutes in the dark at 4°and were washed in FACS Buffer (2% Fetal Bovine Serum in PBS) before analysis on flow cytometer. All the samples were analyzed using BD LSR FORTESSA flow cytometer instrument, which can detect up to 15 different fluorochrome conjugated antibodies simultaneously. An acquisition gate was established based on forward scatter and side scatter parameters that included only monocyte and lymphocyte populations and excluded dead cells and debris.

## Statistical Analysis

Statistical analysis in all plots was performed using R. Statistically significant differences between multiple independent groups in Fig. (2G), (4B) and Supplementary Materials Fig. (3B) were identified by implementing non-parametric Kruskal-Wallis test. Comparisons between the Naïve group at Day 1 and Vaccine at every time point were identified by Post Hoc analysis using Mann Whitney U Test with False Discovery Rate adjustment to control family-wise type I error. Graphs were plotted with GraphPad Prism Version 8.3.0.

In Figs. (3C), (3F), (5A), statistically significant differences between the independent groups at each time point were calculated by non-parametric Mann–Whitney *U* test.

In Figs. (3B), (3E), (5B) statistically significant differences between the independent groups for each organ were defined by non-parametric Mann–Whitney *U* test.

In Figs. (2E) & (6B) statistically significant differences between the independent groups at each time point were identified by unpaired t-test.

Results were deemed as statistically significant when the null hypothesis could be rejected with >95% confidence. Bars represent the mean and standard deviation in all figures.

## Acknowledgments

We thank Dana Bolgen, Arthur Nedder, Kazuo Imaizumi, Sarai Bardales for assistance with mouse and pig models.

## Authors contributions

M.S, E.J.D, M.J.C conceived the project which was directed by D.E.I and D.J.M. Vaccines were prepared and analyzed by B.T.S., D.A.W. A.G.S, N.D., M.K., C.L.H, S.A.R., M.O.D, A.W.L, J.M.S. Experiments in mouse and pig models were conducted by F.L., A.R.G., K.R.L., F.U., C.D.Y., A. J., S.L.L. The data were analyzed by MS., N.L., M.C., N.D. and V. C. The manuscript was written by M.S., E.J.D., D.E.I. and D.J.M. All authors critically reviewed the manuscript.

## Data and materials availability

All data associated with this study are present in the paper or the Supplementary Materials. FcMBL beads can be obtained through an MTA.

## Supplementary Materials

**Supplementary Materials Fig. 1.**
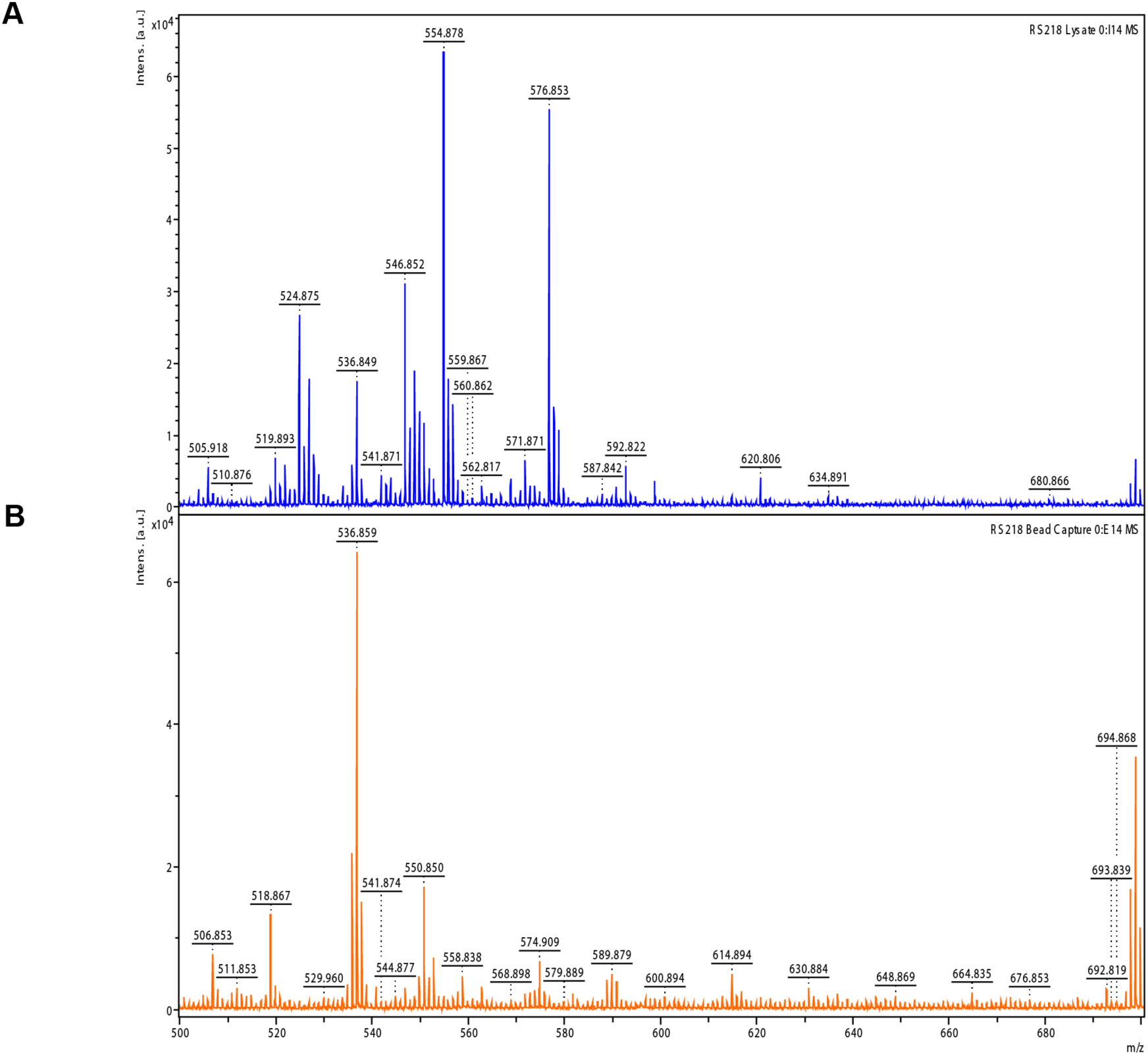
MALDI-TOF Mass Spectrometry differentiates antigens in RS218 lysate from RS218 PAMPs captured on FcMBL beads. **(A)** 1e7 CFU RS218 was killed with ceftriaxone, trypsinized, mixed with TFA and run on mass spectrometry (blue spectrum). Representative profile (blue). **(B)** Ceftriaxone-killed RS218 was mixed with FcMBL-beads, captured PAMPs were enriched by magnetic separation and washing, eluted at 70°C, then trypsinized, mixed with TFA and run on mass spectrometry (brown spectrum). Samples were run in the reflector positive mode below a m/z of 2,000 Da.

**Supplementary Materials Fig. 2.**
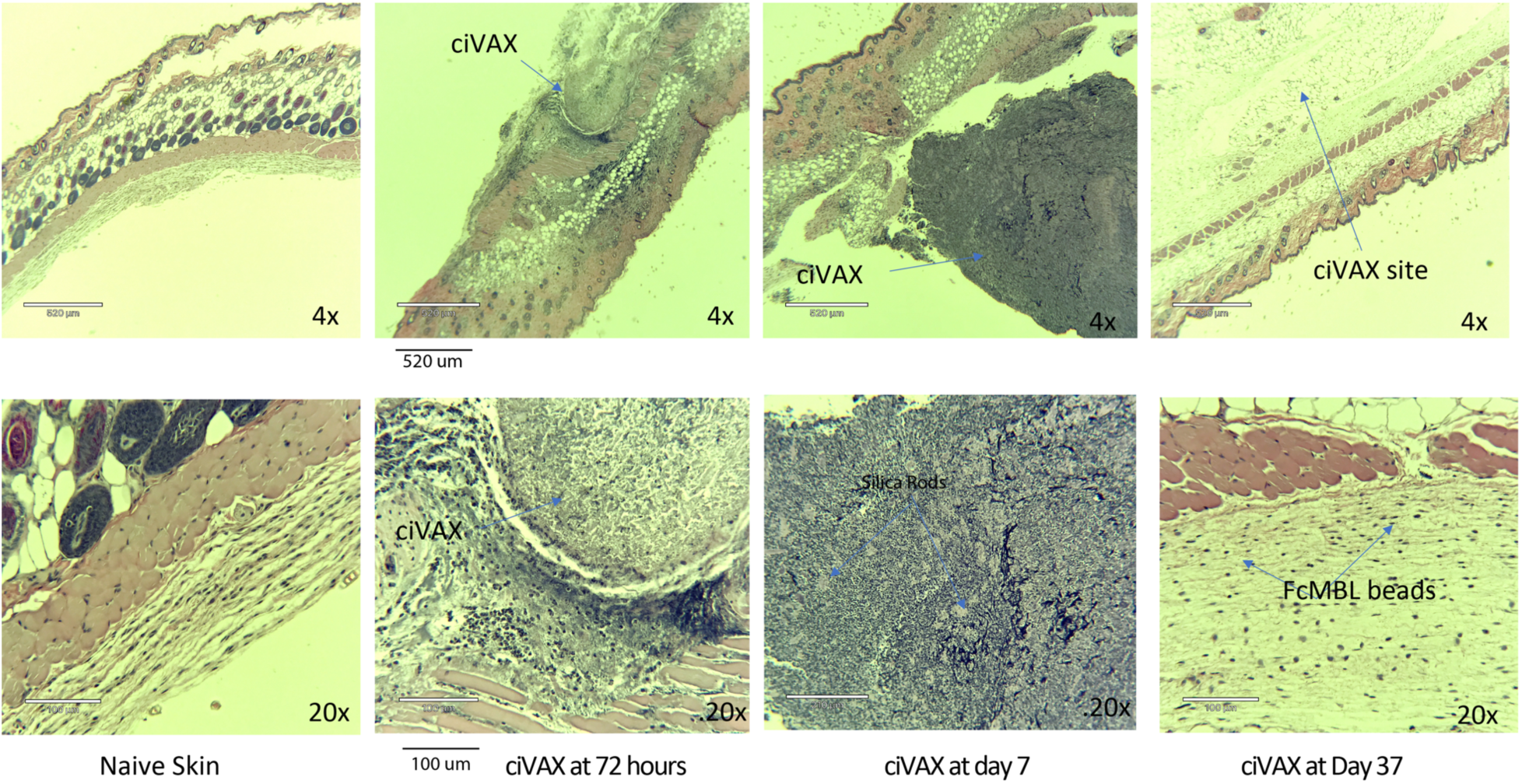
ciVAX produces a long-duration subcutaneous depot. ciVAX was injected by subcutaneous route in mice and compared with naïve mouse skin. The injection sites were dissected at 3, 7 or 37 days. Histology sections were stained with H&E. Scale bars in the upper panels are 520 microns, in the lower panels, these are 100 microns.

**Supplementary Materials Fig. 3.**
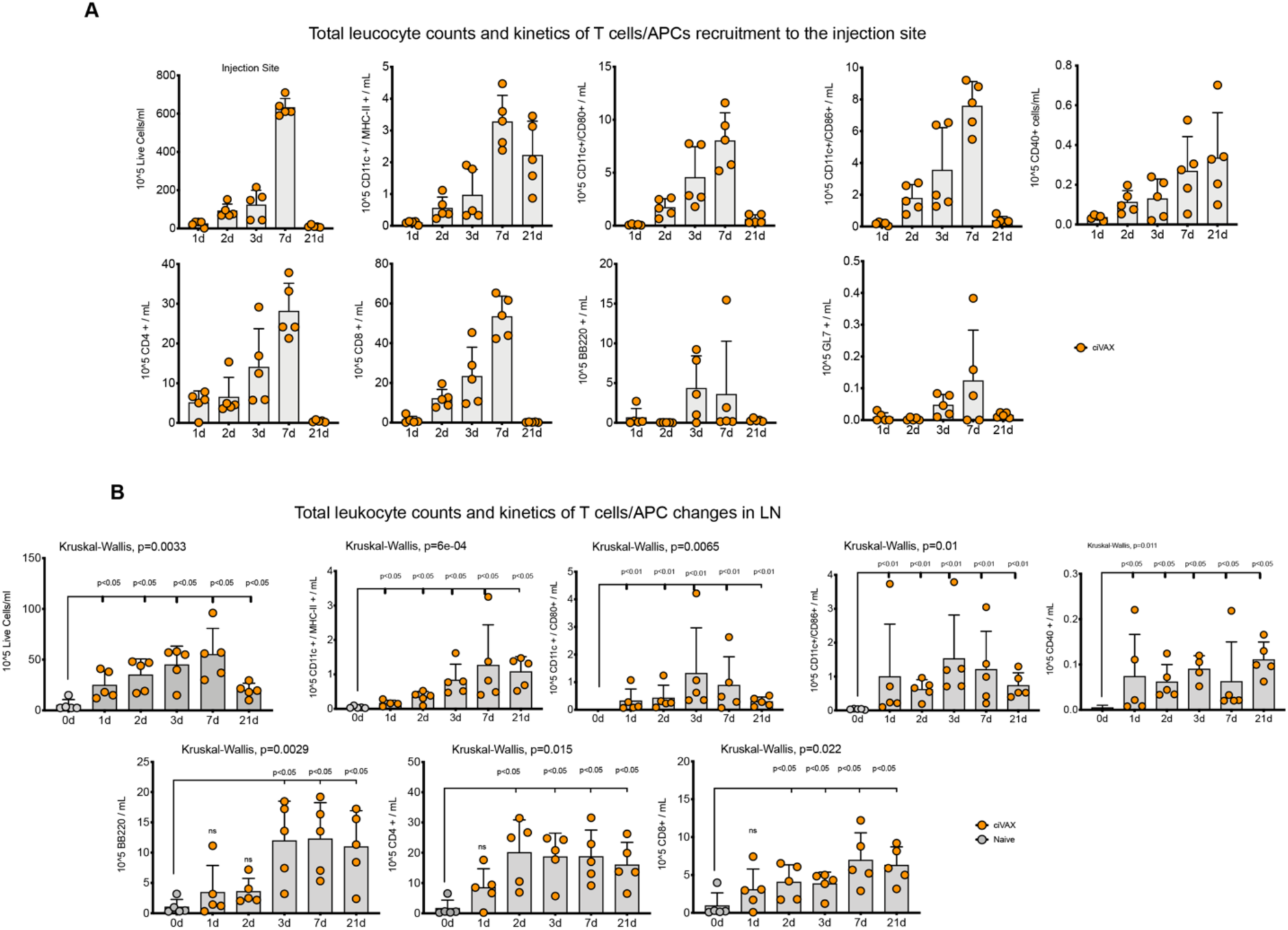
The infection Vaccine induces strong cellular immune responses, with recruitment of DC, B and T cells to the injection site (A), as well as increases in the draining Lymph node (B). Supplementary Materials **Fig.3B**: Statistically significant differences between multiple independent groups were performed using the implementing non-parametric Kruskal-Wallis test. Comparisons between the Naïve group at Day 1 and Vaccine at each time point were identified by Post Hoc analysis using Mann Whitney U Test

**Supplementary Materials Fig. 4 | Minimal changes in microbiome of bacteroides and firmicutes compared with other bacteria after vaccination with the *E. coli* and MRSA ciVAX in rabbits**. (n=2 for each pathogen)

**Supplementary Materials Fig. 4A.**
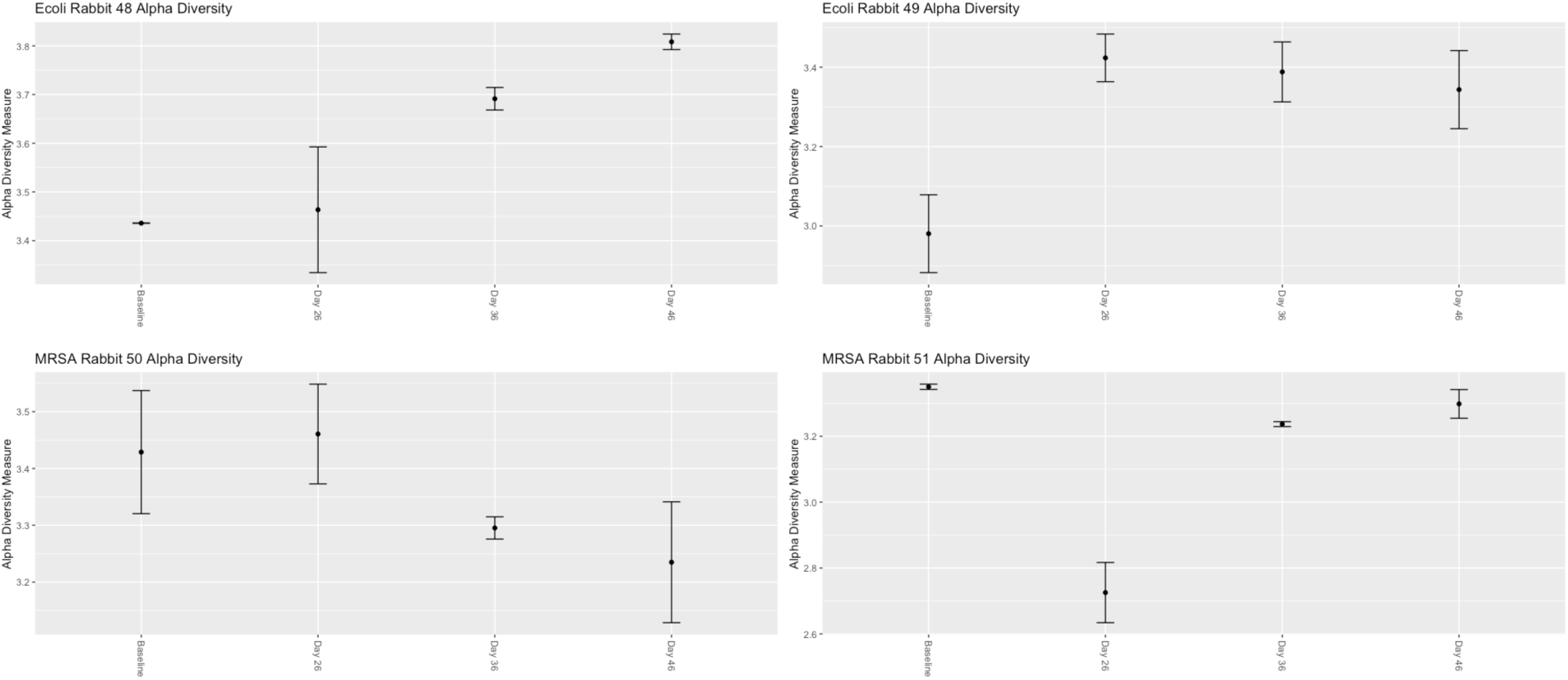
Alpha diversity of the rabbits were measured using Shannon Diversity Index. This index relates both richness and evenness. Where richness is the measure of the number of different species per sample, and evenness is the measure of relative abundance making up the total number of different species. The Shannon Diversity Index was measured at every time point of the experiment.

**Supplementary Materials Fig. 4B.**
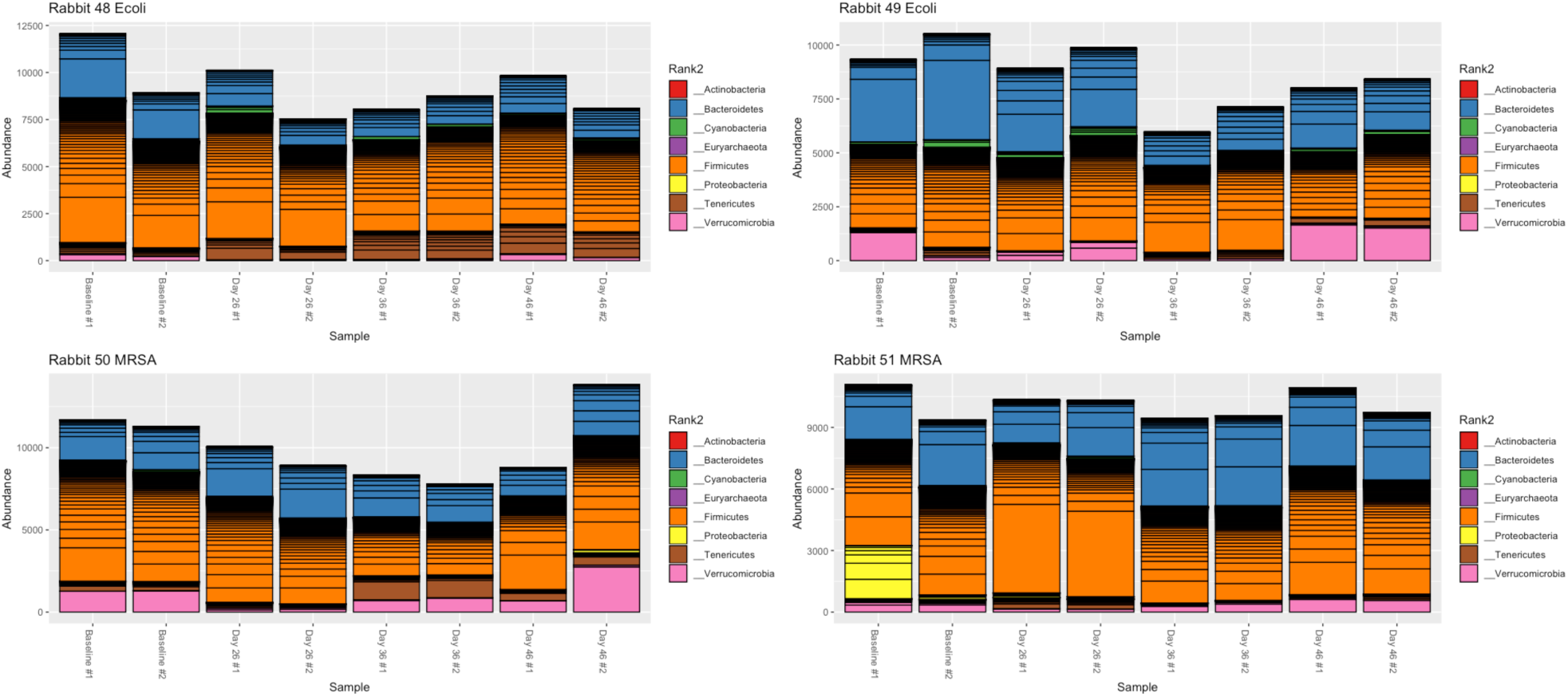
Alpha Diversity was also measured using Taxa abundance table showing timepoints on the x axis and the abundance values on the y axis. For each time point the abundance for each genus is ordered greatest to least and separated by horizontal line. The colors correspond to the respective Phylum.

**Supplementary Materials Fig. 4C.**
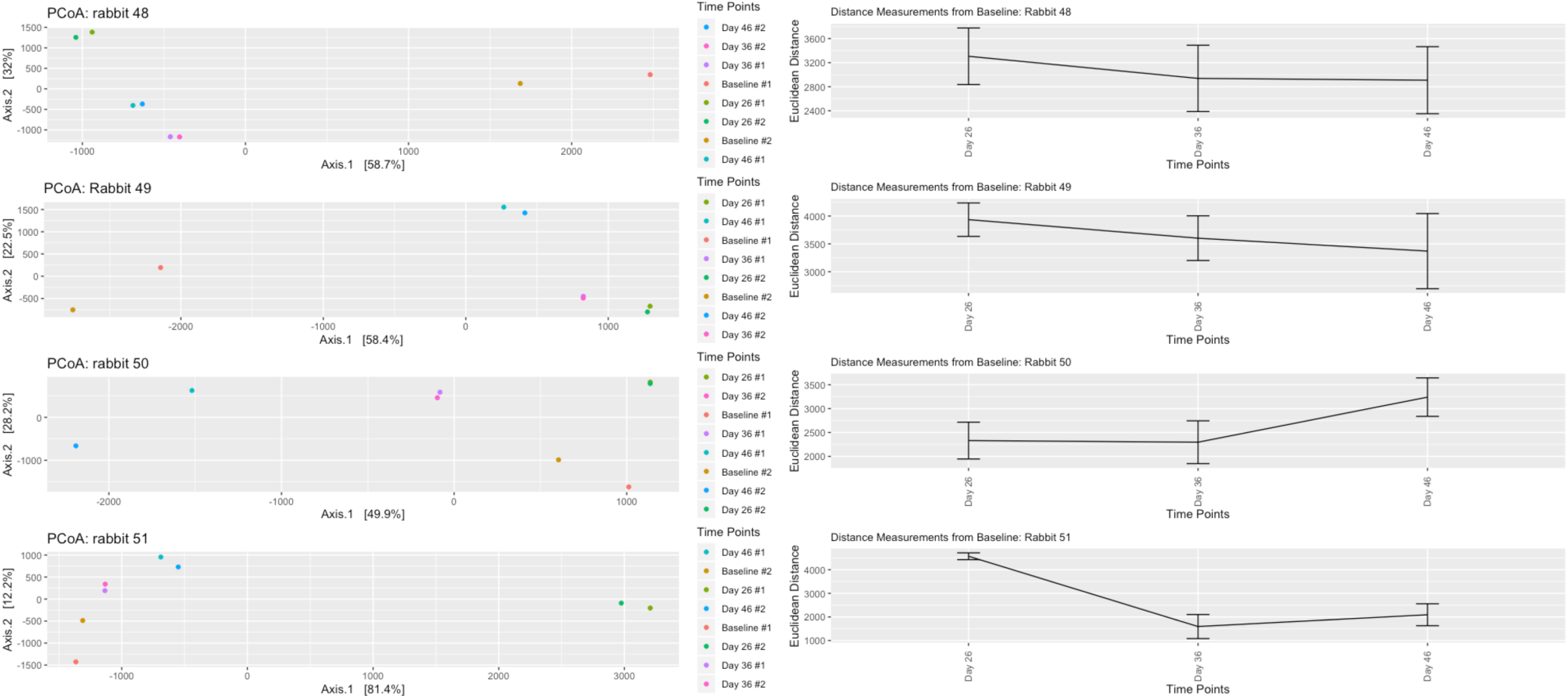
In order to measure beta diversity we performed hierarchal clustering using Principal Component Analysis (PCA). This method is a statistical procedure which uses orthogonal transformations to reduce the number of dimensions in an effort to explain the underlying variance structure of the larger set of variables (i.e. OTU) through a few linear combinations. PCA allows us to determine clustering by using Euclidean distance. Through this analysis we can see how the various time points within samples differ from baseline samples. Since baselines time points have no Vaccine, the relative Euclidean distance between time points and the Baseline, will allow us to determine how different the gut microbiome is after vaccination

**Supplementary Materials Fig. 5.**
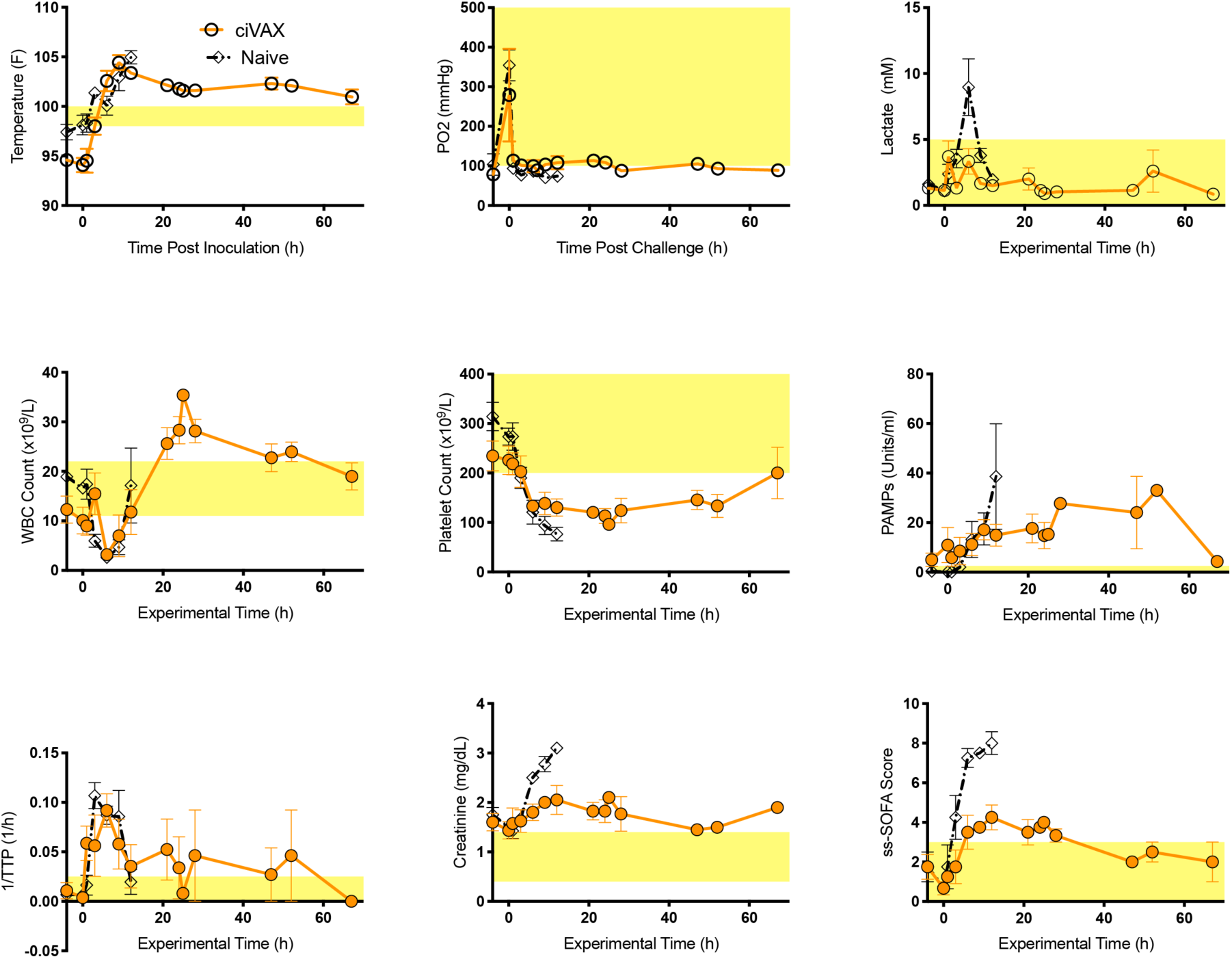
The infection vaccine protects against lethal *E. coli* challenge in a pig model of septic shock. 40-55 kg female juvenile Yorkshire swine (n=4) were vaccinated with ciVAX containing a clinical strain of *E. coli* (isolate 41949 from BWH Crimson Biorepository) and boosted at d28. At d42 they were challenged with (0.5e7-0.5e8 *E. coli* 41949 infused in 4-8 hrs). Normal levels of temperature, pO2, Lactate, WBC, Platelets, PAMPs, 1/TTP, creatinine and swine ssSOFA are highlighted in yellow. Values are not shown for naïve animals (n=4) at later time points as these animals did not survive.

